# CD4^+^ T-cell immunity of SARS-CoV-2 patients determine pneumonia development

**DOI:** 10.1101/2023.05.31.543129

**Authors:** Erika Naito, Takahiro Uchida, Satoshi Yamasaki, Kosuke Hashimoto, Yu Miura, Ayako Shiono, Takayuki Kawamura, Atsuto Mouri, Ou Yamaguchi, Hisao Imai, Kyoichi Kaira, Manabu Nemoto, Kotaro Mitsutake, Makoto Nagata, Hiroshi Kagamu

## Abstract

Most humans infected with SARS-CoV-2 will recover without developing pneumonia. A few SARS-CoV-2 infected patients, however, develop pneumonia, and occasionally develop cytokine storms. In such cases, it is assumed that there is an inadequate immune response to eliminate viral infected cells and an excessive inappropriate immune response causing organ damage, but little is known about this mechanism. In this study, we used single cell RNA sequencing and mass cytometry to analyze peripheral blood T cells from patients hospitalized with proven COVID-19 infection in order to clarify the differences in host immune status among COVID-19 pneumonia cases, non-pneumonia cases, and healthy controls. The results showed that a specific CD4^+^ T cell cluster with chemokine receptor expression patterns, CXCR3^+^CCR4^-^CCR6^+^ (Th1/17), was less abundant in COVID-19 pneumonia patients. Interestingly, these CD4^+^ T-cell clusters were identical to those we have reported to correlate with antitumor immunity and predict programmed cell death (PD)-1 blockade treatment response in lung cancer. The Th1/17 cell percentages had biomarker performance in diagnosing pneumonia cases. In addition, CTLA-4 expression of type17 helper T cells (Th17) and regulatory T cells (Treg) was found to be significantly lower. This indicates that functional suppression of Th17 was less effective and Treg function was impaired in pneumonia cases. These results suggest that imbalance of CD4^+^ T-cell immunity generates excessive immunity that does not lead to viral eradication. This might be a potential therapeutic target mechanism to prevent severe viral infections.

**Importance:** In this observational study, 49 consecutive patients with SARS-CoV-2 infection confirmed by PCR testing and admitted to Saitama Medical University Hospital and Saitama Medical University International Medical Centre between December 4, 2020 and January 17, 2022 were included. Of these 49 patients, 29 were diagnosed with COVID-19 pneumonia by computed tomography (CT) scan (Table 1). The unique CD4^+^ T-cell immunity with less abundant Th1/17 CD4^+^ T-cell cluster and low expression of CTLA-4 in Th17 and Treg was consistently found in SARS-CoV-2 pneumonia patients on admission and 1-week of admission. The imbalance of CD4^+^ T-cell immunity may contribute to develop pneumonia in SARS-CoV-2 virus infected patients by delaying viral clearance and resulting in an excessive immune response.

## Introduction

Most people infected with severe acute respiratory syndrome coronavirus-2 (SARS-CoV-2) that causes the coronavirus disease 2019 (COVID-19) develop mild to moderate illness and may recover without developing pneumonia. However, a few patients develop pneumonia and fatal illness, imposing serious health threats. The SARS-CoV-2 infection triggers the uncontrolled release of cytokines, leading to cytokine storms, which may lead to severe clinical complications. Studies have shown that steroid therapy and treatments to suppress the immune response, such as anti-interleukin (IL)-6 antibody therapy, are effective against COVID-19 pneumonia. These studies suggest that inappropriate, excessive immune responses and failure to eradicate the virus lead to organ damage.^1, 2^ Known clinical risk factors for developing pneumonia include age, obesity, diabetes, and hypertension.^3^ Moreover, genetic abnormalities of toll-like receptor (TLR) 3 and TLR7 involved in type1 interferon (IFN) production and anti-IFN antibodies have been reported as key causes of such immune status; however, the precise underlying mechanism remains unknown.^4^

CD8^+^ T-cells play important roles in clearing virus-infected cells. CD4^+^ T-cells are essential for the priming and clonal expansion of CD8^+^ T-cells; they also mediate the migration of clonally expanded CD8^+^ T-cells to the site of infection and are required for the acquisition of the cell-killing license. CD4^+^ T-cells undergo functional differentiation, known as polarization at priming, to most efficiently eradicate invading microorganisms. Each polarized helper T-cell, such as type 1 helper (Th1), Th2, and Th17, uses distinct effector cells,^5^ such as CD8^+^ T-cells, eosinophils, and neutrophils. If this polarization is not appropriate, inefficient effector cells may be activated, which not only delays the elimination of infectious microorganisms but may also cause organ damage. It has been reported that Th17 with high inflammation-inducing activity is increased in COVID-19 pneumonia and cytokine storm cases and is involved in organ damage.^6–8^ Th1, which uses CD8^+^ T-cells as effectors, provides effective immunity to viral infection; however, recently, it was demonstrated that CXCR3^+^CCR4^−^CCR6^+^ Th1/17, which was originally reported to be important in autoimmune diseases, also plays an important role.^9, 10^ In SARS-CoV-2 infection, nucleocapsid protein-specific CD4^+^ T-cells were reported to be more abundant in Th1 and Th1/17.^11^

Anti-tumor immunity is considered similar to antiviral immunity because anti-tumor T-cells recognize genetic mutational products as neoantigens and attack tumor cells, which is thought to utilize a system that recognizes viral gene products as non-self-antigens. In addition, programmed cell death (PD)-1 inhibitors, widely used in cancer therapy, are based on the application of PD-1 in the phenomenon of T-cell exhaustion shown in a chronic viral infection model. In a recent study using single-cell RNA sequencing (scRNA-seq), we demonstrated that the CD4^+^ T-cell cluster Th7R, comprising CXCR3^+^CCR4^−^CCR6^+^ (Th1/17) and CXCR3^−^CCR4^−^CCR6^+^ (CCR6 SP) CD4^+^ T-cells, but not Th1, represented the strength of antitumor immunity, and predicted the efficacy of PD-1 blockade therapy against lung cancer.^12^ Although both Th1 and Th1/17 expressed Th1-type genes, such as *TBX21*, *CCL4, and IFNγ*, Th1/17 was characterized by expression of *TCF7*, *IL-7 receptor*, and *GZMK,* instead of *PRF1* and *GZMB*.

Therefore, we hypothesized that the T-cell clusters involved in antiviral immunity are diverse, and their balance significantly affects the outcome of viral infections. To test this hypothesis, we analyzed the differentiation and activation/exhaustion status of CD4^+^ and CD8^+^ T-cells in patients with SARS-CoV-2 infection, intending to decipher the differences in T-cell immune status between patients with or without developing pneumonia.

## Methods

### Study Design and Patients

In this observational study, 49 consecutive patients with SARS-CoV-2 infection confirmed by PCR testing and admitted to Saitama Medical University Hospital and Saitama Medical University International Medical Center between December 4, 2020 and January 17, 2022 were included. Of these 49 patients, 29 were diagnosed with COVID-19 pneumonia by computed tomography (CT) scan (**Table 1**). All participants provided written informed consent, and the study protocol was approved by the Internal Review Board of Saitama Medical University International Medical Center (20-066).

**Table 1.**
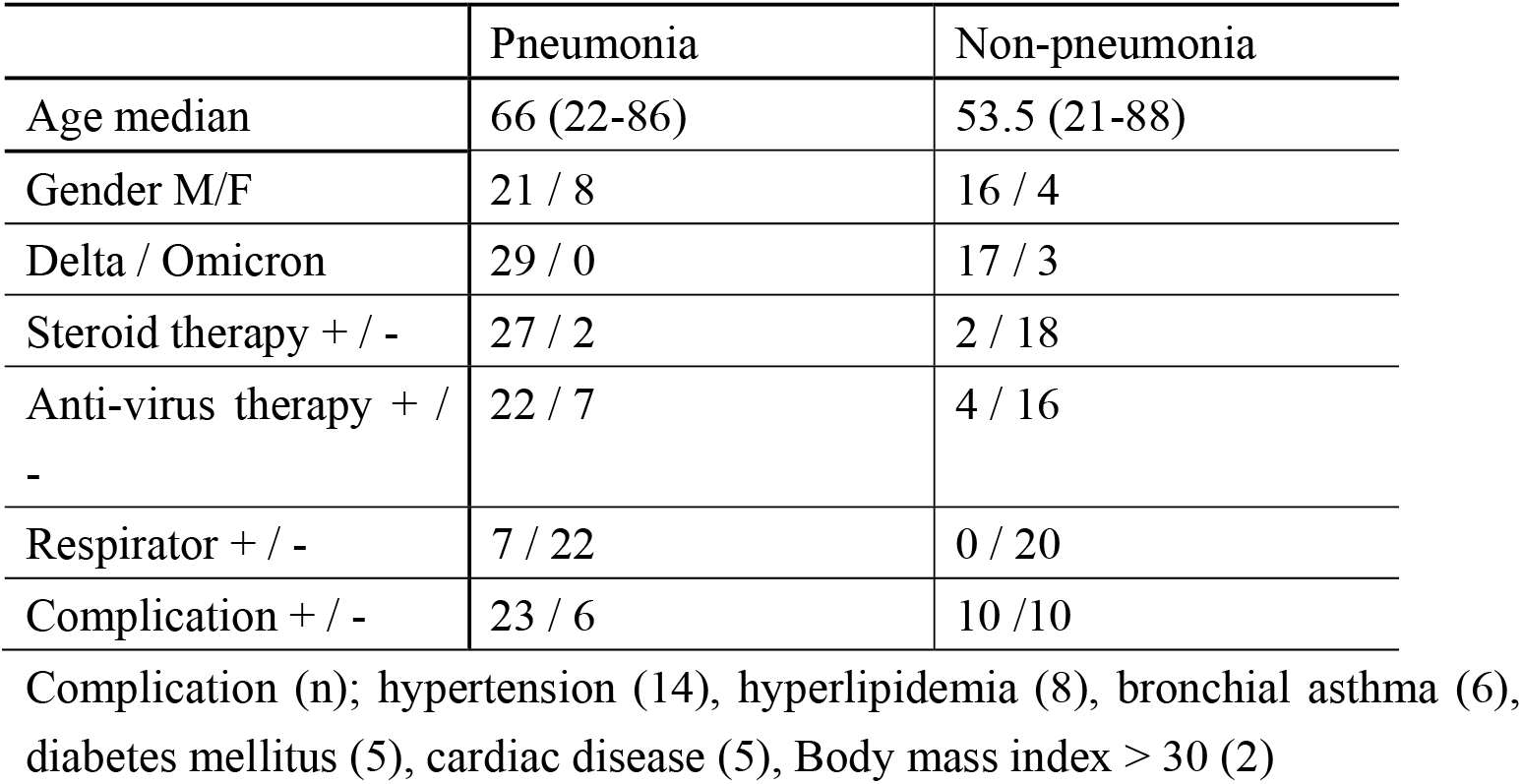
Patient demographics.

|  | Pneumonia | Non-pneumonia |
| --- | --- | --- |
| Age median | 66 (22-86) | 53.5 (21-88) |
| Gender M/F | 21 / 8 | 16 / 4 |
| Delta / Omicron | 29 / 0 | 17 / 3 |
| Steroid therapy + / - | 27 / 2 | 2 / 18 |
| Anti-virus therapy + / - | 22 / 7 | 4 / 16 |
| Respirator + / - | 7 / 22 | 0 / 20 |
| Complication + / - | 23 / 6 | 10 / 10 |

Complication (n); hypertension (14), hyperlipidemia (8), bronchial asthma (6), diabetes mellitus (5), cardiac disease (5), Body mass index > 30 (2)

### Blood Sample Analysis

Blood samples were collected on admission, 1-week after, and on discharge from the hospital, using heparinized CPT Vacutainer tubes (Becton Dickinson Vacutainer Systems, Franklin Lakes, NJ, USA), and peripheral blood mononuclear cells (PBMC) were isolated as described previously.^13^ The PBMC samples were frozen using Cellbanker2 (Nippon Zenyaku Kogyo Co., Koriyama, Japan) in a liquid nitrogen tank. For analyses of T-cell subsets, cells were incubated for 32–48 h in a culture medium RPMI 1640 (Nacalai tesque, Kyoto, Japan) supplemented with 10% fetal calf serum (GIBCO, Carlsbad, CA) before cell staining.

### Mass Cytometry

The monoclonal antibodies used for Helios^TM^ mass cytometry analysis are listed in **Supplementary Table 1 (Table S1)**. Approximately 5.0 × 10^6^ cells were stained with mass cytometry antibodies (Standard BioTools, San Francisco, CA, USA) according to the manufacturer’s instructions. For intracellular staining, samples were prepared using a Maxpar Nuclear Antigen Staining Buffer (Standard BioTools). After washing twice with Maxpar Cell Staining Buffer (Standard BioTools), samples were fixed using Maxpar Fix (Standard BioTools) and Perm Buffer (Standard BioTools) supplemented with 125 nM iridium nucleic acid intercalator (Standard BioTools). Following fixation, cells were resuspended in Maxpar water after washing them with Maxpar Cell Staining Buffer and twice with Maxpar water. More than 2.0 × 10^5^ cells per sample were analyzed using Helios^TM^ and Cytobank (https://www.cytobank.org) software. The gating strategy is presented in **Supplementary Figure 1 (Fig. S1)**.

### Single-cell RNA Sequencing

ScRNA-seq analysis was performed using a total of 10 PBMC samples collected at admission and 1-week later from 5 patients (2 pneumonia and 3 non-pneumonia) using the Chromium Single Cell Immune Profiling v2 kit (10× Genomics Inc., Pleasanton, CA, USA). Details of the scRNA-seq analysis are described in **Supplementary Methods** (**Table S2**).

### Statistical Analysis

Prism 9 (GraphPad Software, San Diego, CA, USA) was used to conduct statistical analyses. Data are expressed as the mean ± standard error of the mean unless otherwise indicated. Tests for differences between two groups were performed using Welch’s *t-*test. Multiple group comparisons were performed using one-way ANOVA followed by Tukey’s *post hoc* analysis. All *P* values were two-sided, and *P* < 0.05 was considered statistically significant.

## Results

### CD4+ T-cell Immunity Differs among Patients With or Without Pneumonia Caused by SARS-CoV-2 Infection and Healthy Controls

ScRNA-seq analysis using the PBMC samples revealed the diversity of T-cell subpopulations at the single-cell gene expression level in patients with (pneumonia group) or without SARS-CoV-2 pneumonia (non-pneumonia group) (**Table S3 and S4**). After the separation of CD4^+^ and CD8^+^ T-cells, we performed unsupervised clustering and uniform manifold approximation and projection (UMAP) analysis of CD4^+^ T-cells based on gene expression of the 2000 features that had highly varied expression levels among all cells (**Figure 1A**). It identified 31 CD4^+^ T-cell clusters, which were annotated based on the gene/protein expression patterns of chemokine receptor proteins and marker genes (CXCR3, CCR4, CCR6, CD45RA, CD62L (*SELL*), and *FoxP3*) (**Figure 1B and Fig. S2A**). The clusters were grouped into four distinct categories: naive CD4^+^ T-cells (comprised CD45RA^+^CD62L^high^; black dashed line in Figure 1B), regulatory T-cells (comprising FoxP3^+^ cells; blue line in Figure 1B), effector T-cells (comprising CD45RA^-^CD62L^low^ CD4^+^ T-cells; red line in Figure 1B), and central memory CD4^+^ T-cells (comprised CD45RA^-^CD62L^high^, others in Figure 1B). To clarify the differences in the number of cells between the groups on the UMAP area, UMAP plane was delimited by grids (grid size = 0.2). The number of cells in the grids was expressed as density (**Figure 1C**). The subtraction of the density in the non-pneumonia group (blue) from those in the pneumonia group (red) on admission identified that the meta-cluster A was more common in the non-pneumonia group, which mainly comprised CD62L^low^CXCR3^+^CCR4^−^CCR6^+^ Th1/17 cell populations (clusters #6 and #23).

**Figure 1.**
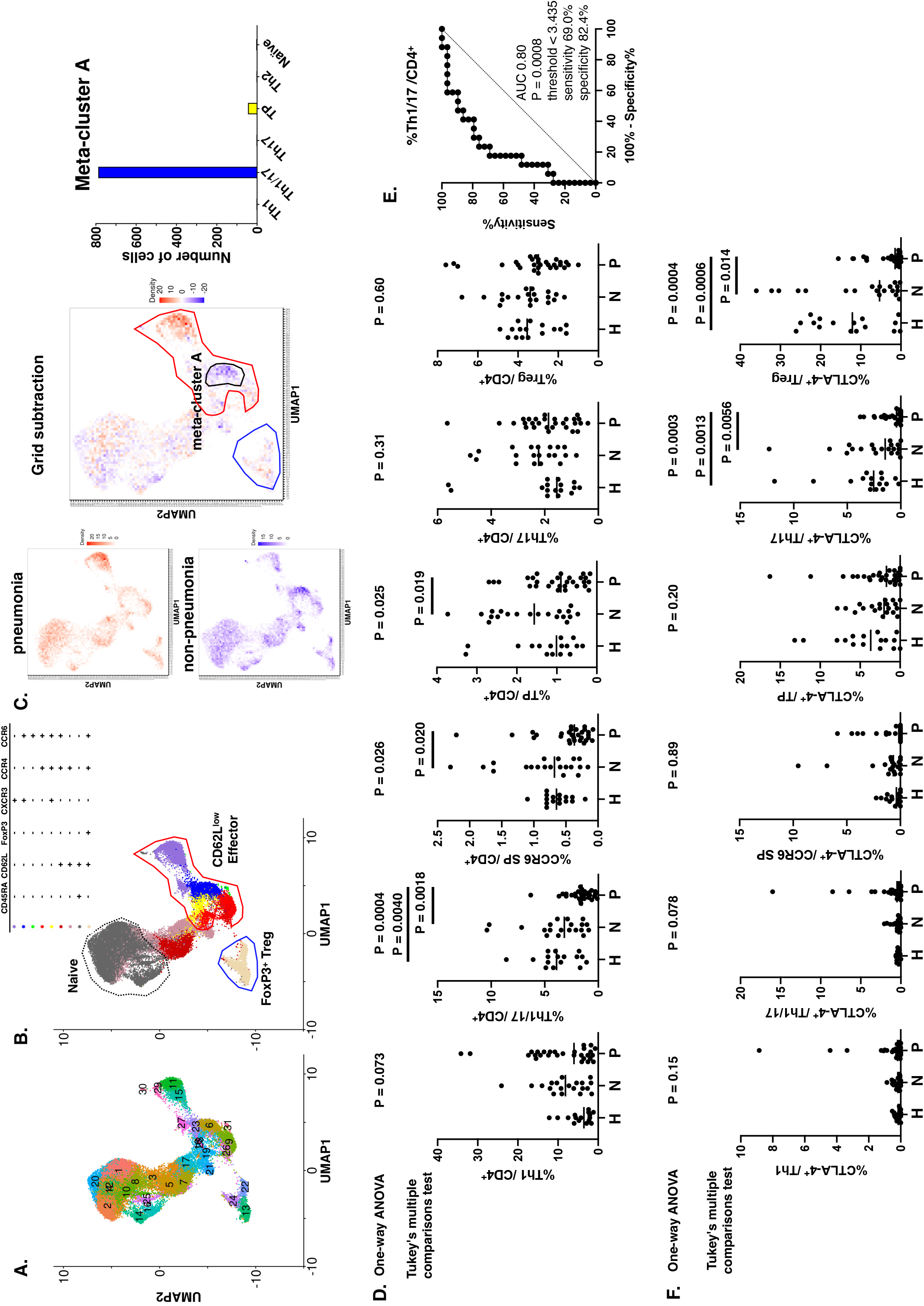
Differences of CD4^+^ T-cell clusters among SARS-CoV-2 pneumonia, non-pneumonia, and healthy control. (**A**) Uniform manifold approximation and projection (UMAP) plots derived from integrated gene expression of CD4^+^ T cells obtained by single-cell RNA sequencing (scRNA-seq) of the samples from patients with SARS-CoV-2 pneumonia (n = 2) and non-pneumonia (n = 3) collected on admission and 1-week after admission. A total of 32,990 CD4^+^ T cells were divided into 31 clusters upon unsupervised clustering. **(B)** CD4^+^ T-cell clusters (n = 31) were annotated according to the gene or protein expression of CD45RA, CD62L, FoxP3, CXCR3, CCR4, and CCR6. The red line indicates the extent of clusters mostly comprising CD62L^low^ effector T cells (clusters #6, 9, 11,15, 18, 19, 23, 27, 28, 29, and 31). The black dashed line indicates clusters #1, 2, 4, 8, 10, 12, 14, 16, 20, and 25—mostly comprising CD62L^high^CD45RA^+^ naïve T-cells. The blue line indicates clusters #13, 22, and 24, enriched with FoxP3^+^ Treg cells. (**C**) The density subtraction of the SARS-CoV-2-infected non-pneumonia group (blue) from those of the SARS-CoV-2-infected pneumonia group (red) at admission. The red line indicates the predominance of patients with pneumonia, the blue line indicates non-the predominance of patients without pneumonia, and the black line indicates meta-cluster A, comprising non-pneumonia patient-predominant cells in effector CD4^+^ T-cell clusters. The bar plot represents the number of cells in meta-cluster A. Meta-cluster A mainly consists of Th1/17 cells (blue dots in Figure 1B). **(D)** The percentages of CD62L^low^CXCR3^+^CCR4^−^CCR6^−^ helper type 1 CD4^+^ T-cell (Th1), CD62L^low^CXCR3^+^CCR4^−^CCR6^+^ Th1/17, CD62L^low^CXCR3^−^CCR4^−^CCR6^+^ CCR6 SP, CD62L^low^CXCR3^+^CCR4^+^CCR6^+^ TP, CD62L^low^CXCR3^−^CCR4^+^CCR6^+^ Th17, and FoxP3^+^ regulatory T-cell (Treg) were compared between patients with SARS-CoV-2 pneumonia (**P**, n = 29), without pneumonia (**N**, n = 20), and healthy control (**H**, n = 15). **(E)** Receiver operating comparison analysis was performed between pneumonia and non-pneumonia groups based on Th1/17 percentages. **(F)** The percentages of CTLA-4^+^ in CD4^+^ T-cell clusters were compared among patients with and without SARS-CoV-2 pneumonia and healthy controls. Statistical analysis was performed using one-way ANOVA, and between-group differences were analyzed using Tukey’s multiple comparisons test.

Next, mass cytometry analysis for determination of CD4^+^ T-cell polarization patterns observed in all patients (n = 49) at admission revealed a significant increase in CD62L^low^CD4^+^ T-cells and Th1 in SARS-CoV-2-infected patients compared to that in healthy controls (**Fig. S2B**). However, the comparison of the three groups revealed a significantly lower percentage of expression of Th1/17 in patients with pneumonia than in those without pneumonia (*P* = 0.0018) and healthy controls (*P* = 0.0040) (**Figure 1D)**; these results are consistent with scRNA-seq results. Furthermore, the percentage of CCR6 single positive (CCR6 SP) and triple positive (TP) CD4^+^ T-cell clusters were significantly lower (*P* = 0.020 and 0.019, respectively) in the patients with pneumonia than those in patients without pneumonia. Receiver Operating Characteristic curve (ROC) analysis showed that patients with pneumonia could be predicted by peripheral Th1/17 with a sensitivity of 69.0% and specificity of 82.4% at a threshold of 3.435 (*P* = 0.0008, **Figure 1E**).

Furthermore, the analysis of CTLA-4 and PD-1 expression in each CD4^+^ T-cell cluster revealed that only Th17 and Treg expressed CTLA-4, and that the pneumonia group had significantly lower CTLA-4 expression in Th17 and Treg (*P* = 0.0056 and 0.014, respectively) than those in the non-pneumonia and healthy control groups (**Figure 1F** and **Fig. S3A, B**). *CTLA-4* gene expression in Tregs was also significantly lower in the pneumonia group than that in the non-pneumonia group (FDR < 0.05, **Fig. S3C**).

### CD8^+^ T-Cell Responses to SARS-CoV-2 Infection

Next, we examined CD8^+^ T-cell clusters separated from CD3^+^ T-cells using scRNA-seq data, as in CD4^+^ T-cells, from unsupervised clustering based on gene expression to grid subtraction (**Figure 2A–C**, **Fig. S4**). CD8^+^ T-cells were classified into 28 clusters by unsupervised clustering, and the annotation of these clusters based on CD45RA and CD62L expression identified an effector memory (EM) cells with CD45RA re-expression (EMRA) region (green line in Figure 2B; comprised CD45RA^+^CD62L^low^CD8^+^ T-cells), an EM region (red line in Figure 2B; comprised CD45RA^−^CD62L^low^), a central memory (CM) T-cell region (yellow line in Figure 2B; comprised CD45RA^−^CD62L^high^), and a naïve CD8^+^ T-cell region (black dashed line in Figure 2B; comprised CD45RA^+^CD62L^high^). The density subtraction in the non-pneumonia group (blue) from the pneumonia group (red) at admission revealed that EM and EMRA CD8^+^ T-cells were more common in patients with pneumonia than those in patients without pneumonia (**Figure 2C**). Mass cytometry analysis showed a lower percentage of naïve and a higher percentage of EMRA in SARS-CoV-2-infected patients than those in healthy controls (**Figure 2D**).

**Figure 2.**
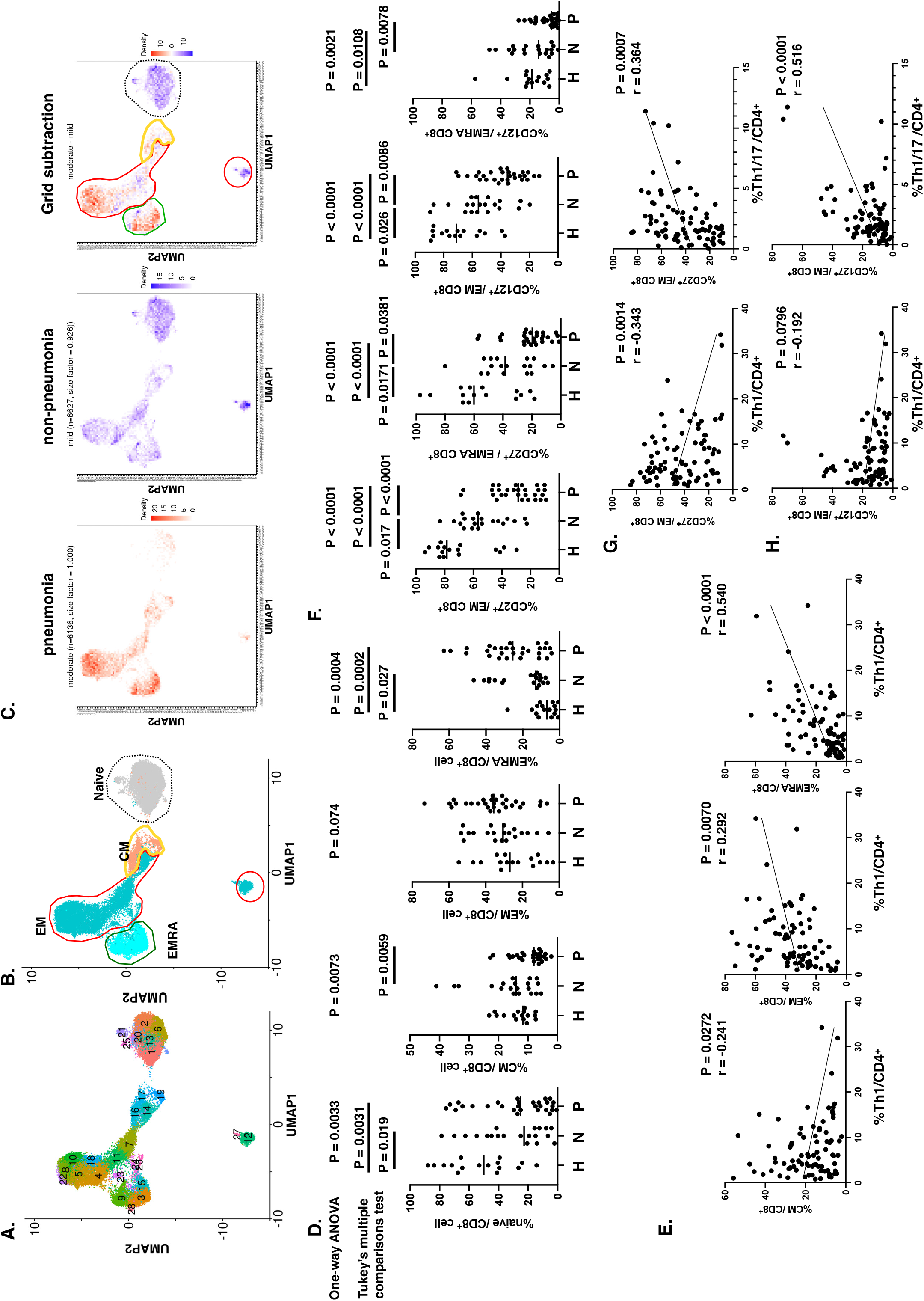
Differences of CD8^+^ T-cell clusters among SARS-CoV-2 pneumonia, non-pneumonia, and healthy control. (**A**) UMAP plots derived from integrated gene expression data of CD8^+^ T-cells from scRNA-seq of two patients with SARS-CoV-2 pneumonia and three patients without SARS-CoV-2 pneumonia collected on admission and 1 week after admission. All 23,236 CD8^+^ T-cells were divided into 28 clusters upon unsupervised clustering. **(B)** CD8^+^ T-cell clusters were annotated according to the gene or protein expression of CD45RA and CD62L (*SELL*). The red line indicates the extent of clusters mainly consisting of CD45RA^-^CD62L^low^ effector memory T-cells (EM, clusters #4, 5, 7, 8, 10, 11, 12, 14, 18, 22, 23, 24, 26, and 27). The green line indicates clusters #3, 9, 15, and 28—mostly comprising CD45RA^+^CD62L^low^ effector memory with re-expression of CD45RA (EMRA). The yellow line indicates clusters #16, 17, and 19, CD45RA^-^CD62L^high^ central memory T-cells (CM). The black dashed line indicates clusters #1, 2, 6, 13, 20, 21, and 25—mostly comprising CD45RA^+^CD62L^high^ naïve T-cells. (**C**) The density subtraction of the SARS-CoV-2-infected non-pneumonia group (blue) from those of the SARS-CoV-2-infected pneumonia group (red) at admission. Red indicates the predominance in patients with pneumonia, and blue indicates the predominance in patients without pneumonia. **(D)** The percentages of EM, EMRA, CM, and naïve T-cell at admission were compared between patients with SARS-CoV-2 pneumonia (**P**, n = 29), non-pneumonia (**N**, n = 20), and healthy control (**H**, n = 15). Statistical analysis was performed using one-way ANOVA, and between-group differences were analyzed using Tukey’s multiple comparisons test. **(E)** Correlations between Th1 and CM, EM, and EMRA. **(F)** The percentages of CD27^+^ and CD127^+^ in CD8^+^ T-cell clusters compared between patients with and without SARS-CoV-2 pneumonia and healthy controls. Statistical analysis was performed using one-way ANOVA, and between-group differences were analyzed using Tukey’s multiple comparisons test. **(G)** Correlations between Th1 and Th1/17 and the percentages of CD27^+^ in EM. **(H)** Correlations between Th1 and Th1/17 and the percentages of CD127^+^ in EM.

To examine the effect of CD4^+^ T-cells on CD8^+^ T-cells that directly eliminate virus-infected cells, we analyzed the correlation between each CD8^+^ T-cell fraction and CD4^+^ T-cell clusters (**Figure 2E**). The analysis identified a positive correlation between Th1 and EMRA. In addition, Th1 showed a weak negative correlation with CM (*P* = 0.0272) and a weak positive correlation with EM (*P* = 0.0070).

Molecular expression analysis of each CD8^+^ T-cell cluster revealed significant differences in the percentages of CD27 and CD127 (IL-7 receptor) expression in EM and EMRA between pneumonia and non-pneumonia groups (**Figure 2F**). Furthermore, the expression of CD27 on EM CD8^+^ T-cells showed a significant positive correlation with Th1/17 but a negative correlation with Th1 in CD4^+^ T-cells (**Figure 2G**). Th1/17 was also significantly positively correlated with IL-7 receptor expression on EM CD8^+^ T-cells (**Figure 2H**).

### CD8^+^ T-cell Immunity Returned to a Steady State during Recovery from SARS-CoV-2 Infection

After admission, 2 of 20 patients without pneumonia and 27 of 29 with pneumonia received steroid therapy. Seven patients with pneumonia received ventilator management, and all recovered eventually. Analysis of PBMC samples one week after hospitalization demonstrated the changes in CD8^+^ T-cell immunity during the recovery phase. The results showed a significant decrease in EMRA fraction. The reduction in the EMRA fraction was proportional to the increase in the CM fraction (**Figure 3A–C**). This result indicates that CD8^+^ T-cell differentiation, which was induced in the EMRA fraction by the viral infection, changed to the CM fraction upon recovery. The decrease in CD27 and CD127 in the effector CD8^+^ T-cell fraction, which was stronger in the pneumonia group than that in the non-pneumonia group, also recovered, and no significant difference was observed in the CD27^+^ and CD127^+^ cells between pneumonia and non-pneumonia groups (**Figure 3D, E**). As described above, the difference in CD8^+^ T-cell fraction between pneumonia and non-pneumonia groups vanished with recovery from SARS-CoV-2 infection.

**Figure 3.**
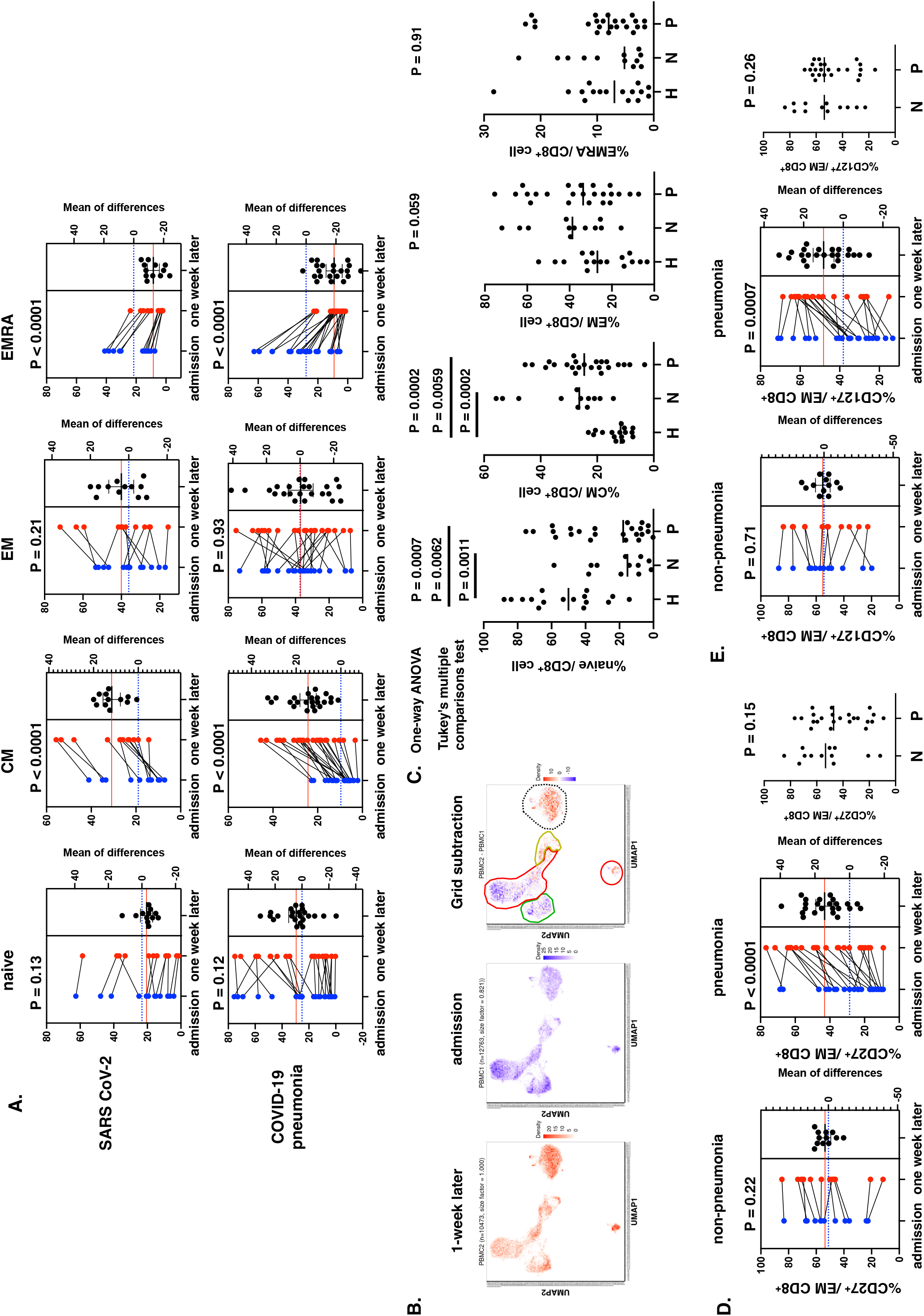
Changes of CD8^+^ T-cell clusters one week after admission. (**A**) Correlation analysis between CD8^+^ T-cell clusters at admission and one week after admission in SARS-CoV-2 pneumonia and non-pneumonia patients (n = 35). **(B)** The subtraction of density of cells at admission (blue) from those one week after admission (red) of five patients. Red indicates predominance one week after, and blue indicates predominance at admission. The red line indicates effector memory T-cells (EM). The green line indicates effector memory with re-expression of CD45RA (EMRA). The yellow line indicates central memory T-cells (CM). The black dashed line indicates naïve T-cells. **(C)** The percentages of EM, EMRA, CM, and naïve CD8^+^ T-cell one week after admission were compared among SARS-CoV-2 pneumonia patients (**P**, n = 23), non-pneumonia patients (**N**, n = 12), and healthy control (**H**, n = 15). Statistical analysis was performed using one-way ANOVA, and between-group differences were analyzed using Tukey’s multiple comparisons test. **(D, E)** Correlation analysis between CD27^+^ and CD127^+^ CD8^+^ T-cell percentages at admission and one week later in SARS-CoV-2 pneumonia and non-pneumonia patients (n = 35). The percentages of CD27^+^ and CD127^+^ CD8^+^ T-cell one week after admission were compared between SARS-CoV-2 pneumonia and non-pneumonia groups.

### CD4^+^ T-cell Immunity Differs Between Patients With or Without Pneumonia After Recovery from SARS-CoV-2 Infection

Analysis of the changes in the CD4^+^ T-cell cluster during recovery from SARS-CoV-2 infection revealed that Th1 decreased significantly in both pneumonia (*P* = 0.0006) and non-pneumonia groups (*P* = 0.0043) one week after admission compared to that at the time of admission (**Figure 4A**). In contrast, Th17 increased; the extent of the increase was greater in the pneumonia group than that in the non-pneumonia group (**Figure 4B**). Furthermore, unlike CD8^+^ T-cell clusters, Th1/17 showed consistent differences between pneumonia and non-pneumonia groups at one week of admission (*P* = 0.0021, *P* = 0.039, respectively; **Figure 4C**). These are consistent with the results of scRNA-seq analysis (**Figure 4D**). The subtraction of the density in the non-pneumonia group (blue) from the pneumonia group (red) at one-week of admission identified meta-cluster B, in which Th1/17 was a major component, was less common in the pneumonia group (**Figure 4C**), as it was at admission (**Figure 1C**).

**Figure 4.**
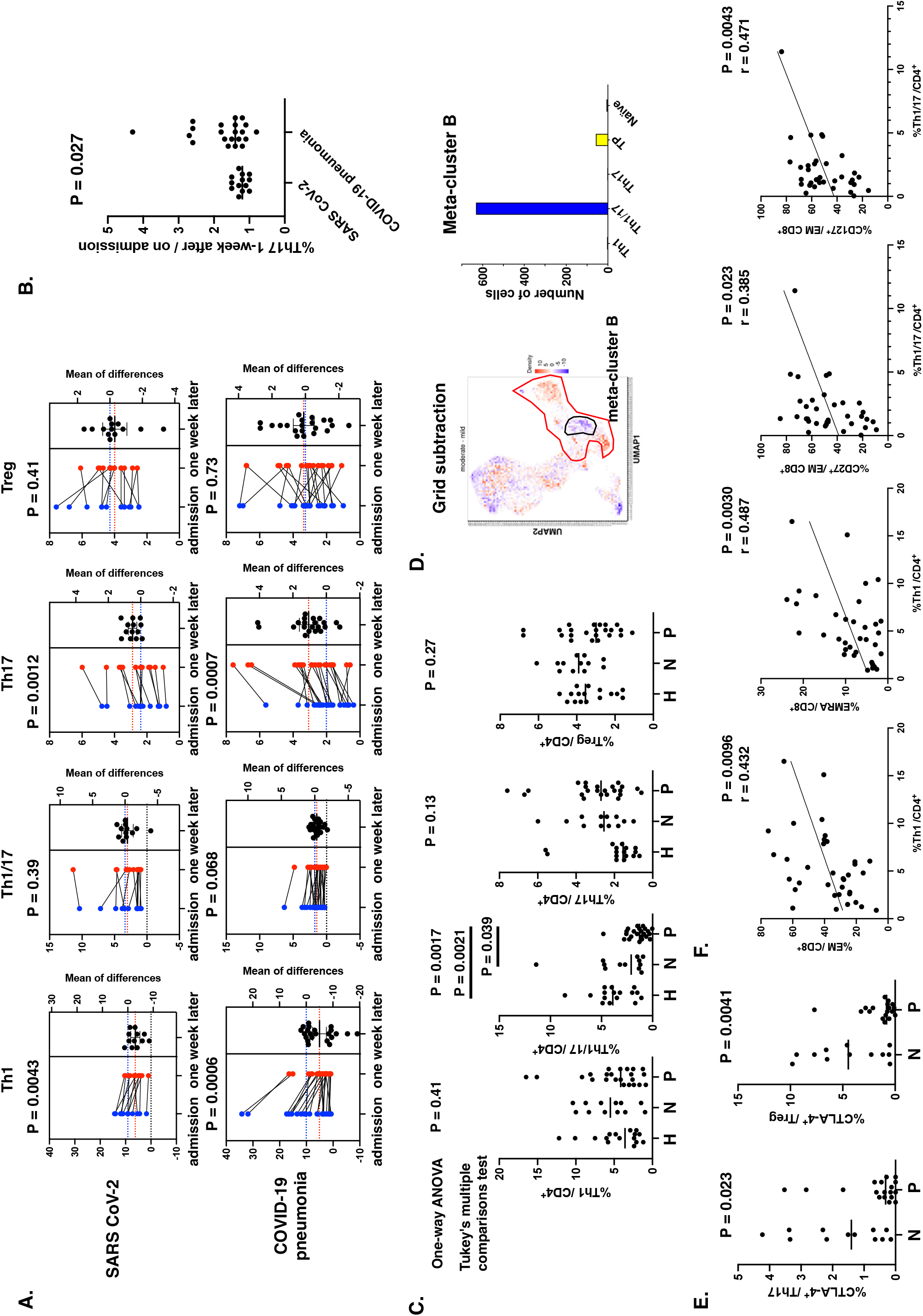
Changes of CD4^+^ T-cell clusters at one week after admission. (**A**) Correlation analysis between CD4^+^ T-cell clusters (Th1, Th1/17, Th17, and Treg) at admission and one week later in SARS-CoV-2 pneumonia and non-pneumonia patients (n = 35). (**B**) The ratio of CD62L^low^ Th17 one week after admission to that at the admission of patients with SARS-CoV-2 pneumonia and non-pneumonia patients was compared. **(C)** The percentages of Th1, Th1/17, Th17, and Treg, one week after admission, compared among SARS-CoV-2 pneumonia patients (**P**, n = 23), non-pneumonia patients (**N**, n = 12), and healthy control (**H**, n = 15). Statistical analysis was performed by one-way ANOVA, and between-group differences were analyzed by Tukey’s multiple comparisons test. **(D)** The density subtraction of the SARS-CoV-2-infected non-pneumonia group (blue) from those of the SARS-CoV-2-infected pneumonia group (red) at one week after admission. Red indicates pneumonia patient-predominant, and blue indicates non-pneumonia patient-predominant cells. The red line indicates the extent of clusters mostly composed of CD62L^low^ effector T-cells. The black line indicates meta-cluster B, consisting of non-pneumonia patient-predominant cells in effector CD4^+^ T-cell clusters. The location of meta-cluster B on the UMAP is almost identical to that of meta-cluster A (Figure 1C), and then meta-cluster B mainly consists of Th1/17 cells. **(E)** The percentages of CTLA-4^+^ in Th17 and Treg one week after admission were compared between patients with (n = 23) and without (n =15) SARS-CoV-2 pneumonia. **(F)** The correlation between the percentage of CD27^+^ and CD127^+^ in EM or EMRA and Th1 or Th1/17 one week after admission was analyzed.

The difference in CTLA-4 positivity between Th17 and Tregs, as indicated by PBMC analysis at admission, persisted after 1 week and was lower in the pneumonia group (*P* = 0.023, *P* = 0.0041, respectively) than that in the non-pneumonia group (**Figure 4E**). The observed correlation between CD8^+^ T-cell clusters, CD27^+^, and CD127^+^ based on effector CD8^+^ T-cells and CD4^+^ T-cell clusters was maintained after 1 week—Th1 was positively correlated with CD8+ T-cell EM and EMRA proportions and Th1/17 was positively correlated with CD27^+^ and CD127^+^ percentages (**Figure 4F**).

## Discussion

Analysis of PBMC T-cells in this study revealed significant differences in T-cell immunity between patients who were infected with SARS-CoV-2 but did not develop pneumonia and those who developed pneumonia.

On admission, the EMRA percentage of CD8^+^ T-cells was higher in patients with SARS-CoV-2 infection, regardless of whether they had pneumonia; however, the percentage decreased to the same level as that in healthy controls after one week of admission. The increase in CM instead of EMRA suggested that effector CD8^+^ T-cells, which had increased in response to SARS-CoV-2 infection, became central memory as the viral load decreased. In contrast to this dynamic change in CD8^+^ T-cell clusters, the differences found in CD4^+^ T-cell clusters between pneumonia and non-pneumonia groups in Th1/17 percentage and CTLA-4^+^ Th17 and Tregs were consistent. The percentages of Th1/17 in CD4^+^ T-cell, CTLA-4^+^ Th17, and CTLA-4^+^ Tregs were significantly lower in the pneumonia group than those in the non-pneumonia group. This study also revealed that Th1 and Th1/17, which play crucial roles in viral immunity, have different helper functions for CD8^+^ T-cells. The positive correlation of Th1 with EMRA CD8^+^ T-cells and negative correlation with CD27 expression in effector CD8^+^ T-cells indicated its role in promoting terminal differentiation of CD8^+^ T-cells with high cytotoxic activity and low co-stimulatory signaling through CD27.^6, 14^ In contrast, the positive correlation of Th1/17 with CD27^+^ and CD127^+^ in effector CD8^+^ T-cells suggested its role in homeostatic lymphocyte proliferation, and effector T-cell development, maintenance, and survival.^10, 14–17^ Furthermore, long-lived memory T-cells in SARS-CoV-2-specific effector CD8^+^ T-cells have been shown to express IL-7 receptors.^18^ Therefore, it is possible that in patients without pneumonia, the ability of CD8^+^ T-cells to kill and maintain survival is balanced. In contrast, in patients with SARS-CoV-2 pneumonia, only the killing function of CD8^+^ T-cells is prominent, while CD8^+^ T-cell survival ability was reduced, possibly affecting their ability to eliminate virus-infected cells. These results suggest that the differences in Th1/17 but not Th1 determine susceptibility to SARS-CoV-2 pneumonia. This result is consistent with that of our previous study, which reported the efficacy of the effective T-cell clusters for PD-1 inhibitors against lung cancer.^12^ Our previous study also showed that Th1/17, but not Th1, exhibits anti-tumor T-cell immune potency.^12^ Taken together, we inferred that the same CD4^+^ T-cell cluster that plays an important role in anti-tumor immunity by recognizing gene mutation products presented on tumor cells also plays a key role in viral immunity by recognizing viral gene products on virus-infected cells. Furthermore, we demonstrated that during recovery from the viral infection, the differences in CD4^+^ T-cell clusters were consistent, unlike those in CD8^+^ T-cells. This result suggests that differences in CD4^+^ T-cell immunity reflect individual host immunity differences.

Th17, with a high inflammatory potential, which uses neutrophils as effector cells, causes acute respiratory distress syndrome.^19^ Increased Th17/Treg ratio is also associated with LPS-induced acute lung injury and respiratory syncytial virus infection.^19, 20^ In addition, Th17 plays an important role in developing pneumonia in patients with SARS-CoV-2-infection. In the present study, we found that CTLA-4 expression of Th17 and Tregs was significantly decreased in patients with pneumonia, which suggests that the important inhibitory mechanism of Th17 is reduced, making Th17 more likely to be activated and Treg function impaired in the pneumonia group. The percentages of CTLA-4^+^ on Th17 and Treg were also not significantly altered in patients who recovered from the viral infection, suggesting that the low percentages of CTLA-4 on Th17 and Treg are a host characteristic rather than a transient response to virus infection.

One limitation of this study is that the antigen specificity of T-cells was not analyzed. However, previous reports have shown that SARS CoV-2 membrane antigen-specific and nucleocapsid antigen-specific CD4^+^ T-cells had preferential polarization to Th1 or Th1/17. This suggests that a balance of CD4^+^ T-cells with different help functions for the same antigen is established. Further analysis is needed to determine whether the differences of Th1/17 and CTLA-4 expression on Th17 and Treg are constitutional, supported by SNPs or epigenesis, and their pathogenic roles in virus infections.

This study showed that the imbalance of CD4^+^ T-cell immunity is an important factor in the pathogenesis of pneumonia caused by SARS-CoV-2 infection. Moreover, it suggests that peripheral Th1/17 is a biomarker for SARS-CoV-2 pneumonia susceptibility. The findings of this study provide insights into the role of host immune status in identifying patients at high risk for severe viral infections and determining their responses appropriately, which may assist in avoiding severe cases due to pandemics by minimizing therapeutic intervention.

## Data Sharing Statement

The RAW data of scRNA-seq generated in this study are publicly available in Gene Expression Omnibus (GEO) at 224482. The analytical codes of scRNA-seq analysis are publicly available in Code Ocean (https://codeocean.com/capsule/1754021/).

## Additional Contributions

We thank Dr. Katsuhisa Horimoto for critical discussion.

## References

1. Zhang Q, Bastard P, Effort CHG, Cobat A, Casanova JL. Human genetic and immunological determinants of critical COVID-19 pneumonia. Nature. Mar 2022;603(7902):587–598. doi:10.1038/s41586-022-04447-0

2. Tay MZ, Poh CM, Renia L, MacAry PA, Ng LFP. The trinity of COVID-19: immunity, inflammation and intervention. Nat Rev Immunol. Jun 2020;20(6):363–374. doi:10.1038/s41577-020-0311-8

3. Grasselli G, Zangrillo A, Zanella A, et al. Baseline Characteristics and Outcomes of 1591 Patients Infected With SARS-CoV-2 Admitted to ICUs of the Lombardy Region, Italy. JAMA. Apr 28 2020;323(16):1574–1581. doi:10.1001/jama.2020.5394

4. Zheng HY, Zhang M, Yang CX, et al. Elevated exhaustion levels and reduced functional diversity of T cells in peripheral blood may predict severe progression in COVID-19 patients. Cell Mol Immunol. May 2020;17(5):541–543. doi:10.1038/s41423-020-0401-3

5. Sallusto F. Heterogeneity of Human CD4(+) T Cells Against Microbes. Annu Rev Immunol. May 20 2016;34:317–34. doi:10.1146/annurev-immunol-032414-112056

6. Xu Z, Shi L, Wang Y, et al. Pathological findings of COVID-19 associated with acute respiratory distress syndrome. Lancet Respir Med. Apr 2020;8(4):420–422. doi:10.1016/S2213-2600(20)30076-X

7. Wu D, Yang XO. TH17 responses in cytokine storm of COVID-19: An emerging target of JAK2 inhibitor Fedratinib. J Microbiol Immunol Infect. Jun 2020;53(3):368–370. doi:10.1016/j.jmii.2020.03.005

8. Orlov M, Wander PL, Morrell ED, Mikacenic C, Wurfel MM. A Case for Targeting Th17 Cells and IL-17A in SARS-CoV-2 Infections. J Immunol. Aug 15 2020;205(4):892–898. doi:10.4049/jimmunol.2000554

9. Manh DH, Weiss LN, Thuong NV, et al. Kinetics of CD4(+) T Helper and CD8(+) Effector T Cell Responses in Acute Dengue Patients. Front Immunol. 2020;11:1980. doi:10.3389/fimmu.2020.01980

10. Chng MHY, Lim MQ, Rouers A, et al. Large-Scale HLA Tetramer Tracking of T Cells during Dengue Infection Reveals Broad Acute Activation and Differentiation into Two Memory Cell Fates. Immunity. Dec 17 2019;51(6):1119–1135 e5. doi:10.1016/j.immuni.2019.10.007

11. Sekine T, Perez-Potti A, Rivera-Ballesteros O, et al. Robust T Cell Immunity in Convalescent Individuals with Asymptomatic or Mild COVID-19. Cell. Oct 1 2020;183(1):158–168 e14. doi:10.1016/j.cell.2020.08.017

12. Kagamu H, Yamasaki S, Kitano S, et al. Single-Cell Analysis Reveals a CD4+ T-cell Cluster That Correlates with PD-1 Blockade Efficacy. Cancer Res. Dec 16 2022;82(24):4641–4653. doi:10.1158/0008-5472.CAN-22-0112

13. Kagamu H, Kitano S, Yamaguchi O, et al. CD4(+) T-cell Immunity in the Peripheral Blood Correlates with Response to Anti-PD-1 Therapy. Cancer Immunol Res. Mar 2020;8(3):334–344. doi:10.1158/2326-6066.CIR-19-0574

14. Powell DJ, Jr., Dudley ME, Robbins PF, Rosenberg SA. Transition of late-stage effector T cells to CD27+ CD28+ tumor-reactive effector memory T cells in humans after adoptive cell transfer therapy. Blood. Jan 1 2005;105(1):241–50. doi:10.1182/blood-2004-06-2482

15. Tsunetsugu-Yokota Y, Kobayahi-Ishihara M, Wada Y, et al. Homeostatically Maintained Resting Naive CD4(+) T Cells Resist Latent HIV Reactivation. Front Microbiol. 2016;7:1944. doi:10.3389/fmicb.2016.01944

16. Shi LZ, Fu T, Guan B, et al. Interdependent IL-7 and IFN-gamma signalling in T-cell controls tumour eradication by combined alpha-CTLA-4+alpha-PD-1 therapy. Nat Commun. Aug 8 2016;7:12335. doi:10.1038/ncomms12335

17. Xu H, Bendersky VA, Brennan TV, Espinosa JR, Kirk AD. IL-7 receptor heterogeneity as a mechanism for repertoire change during postdepletional homeostatic proliferation and its relation to costimulation blockade-resistant rejection. Am J Transplant. Mar 2018;18(3):720–730. doi:10.1111/ajt.14589

18. Adamo S, Michler J, Zurbuchen Y, et al. Signature of long-lived memory CD8(+) T cells in acute SARS-CoV-2 infection. Nature. Feb 2022;602(7895):148–155. doi:10.1038/s41586-021-04280-x

19. Li JT, Melton AC, Su G, et al. Unexpected Role for Adaptive alphabetaTh17 Cells in Acute Respiratory Distress Syndrome. J Immunol. Jul 1 2015;195(1):87–95. doi:10.4049/jimmunol.1500054

20. Mangodt TC, Van Herck MA, Nullens S, et al. The role of Th17 and Treg responses in the pathogenesis of RSV infection. Pediatr Res. Nov 2015;78(5):483–91. doi:10.1038/pr.2015.143

